# Female-biased continental farmer ancestry contribution to the modern Japanese genome

**DOI:** 10.64898/2026.07.28.741206

**Authors:** Yusuke Watanabe, Yoshiki Wakiyama, Jun Ohashi, NCBN Controls WGS Consortium, Hiroki Oota

## Abstract

Modern mainland Japanese formed through admixture between the Jomon hunter– gatherers and continental farmers who arrived probably in northern Kyushu around 3,000 years ago (the dual-structure model). Although Jomon ancestry accounts for only ∼10–20% of the autosomes, the Jomon-associated Y-chromosome haplogroup D1a2a reaches ∼35% in present-day Japanese males—an “admixture paradox” often attributed to a larger contribution by Jomon males, though this has never been tested genome-wide. To test whether this admixture was sex-biased, here we compared the X chromosome with the autosomes. Using Jomon-derived variants as local-ancestry markers, we separated the modern Japanese genome into Jomon-related and continental East Asian-related ancestry components and computed the Keinan-type autosome-to-X drift ratio (Q, neutral expectation 0.75) for each. The continental East Asian-related component gave a significantly higher Q than the Jomon-related component (Δ = Q_farming_ − Q_HG_ = 0.10, P = 9.1 × 10 □³). Relative to the neutral value of 0.75, the Jomon-related component was indistinguishable from neutrality (Q = 0.78, P = 0.34), whereas the continental East Asian-related component was significantly higher (Q = 0.88, P = 4.2 × 10□ □). Under a single-pulse founder model, this pattern implies a continental farmer contribution that was significantly more female-biased than the Jomon hunter-gatherer contribution, with no detectable departure from sex balance in the latter. Monte Carlo simulations showed that the inferred sex ratios account for much of the observed D1a2a frequency, although the remaining discrepancy indicates that additional demographic processes may have contributed. The ancestry asymmetry in the modern Japanese genome was thus driven primarily by a female-biased contribution of continental farming migrants, rather than by the male-biased Jomon contribution conventionally invoked to explain the admixture paradox.

## Introduction

The formation of modern Japanese has been understood mainly through the dual-structure model (Hanihara, 1991), in which the present-day populations arose through admixture between the Jomon hunter–gatherers and continental farmers (who immigrated around 3,000 years ago, which by definition corresponds to the beginning of the Yayoi period). This model proposed by morphological studies has been broadly supported by later genetic studies, first by classical markers (Omoto and Saitou, 1997) and later by genome-wide SNP data and ancient genomes (Yamaguchi-Kabata et al., 2008; Japanese Archipelago Human Population Genetics Consortium, 2012; Jinam et al., 2015; Nakagome et al., 2015; Kanzawa-Kiriyama et al., 2017, 2019; Gakuhari et al., 2020; Cooke et al., 2021; Watanabe et al., 2021; Watanabe and Ohashi, 2023; Liu et al., 2024; Yamamoto et al., 2024). The Jomon are a deeply diverged, basal East Asian lineage (Kanzawa-Kiriyama et al., 2019; Gakuhari et al., 2020; Cooke et al., 2021), and their ancestry accounts for roughly 10–20% of the autosomes of present-day mainland Japanese (Kanzawa-Kiriyama et al., 2017, 2019; Gakuhari et al., 2020; Cooke et al., 2021).

Prehistoric changes in subsistence and movements of people were often sex-specific. When two populations mix, the two sexes can contribute unequally: kinship systems, marriage and residence customs, and sex differences in mobility all shape who ends up transmitting their genes. Because the paternally inherited Y chromosome and the maternally inherited mtDNA each trace a single sex, whereas the autosomes average over both, such imbalances leave a distinctive genetic signature (Seielstad et al., 1998; Oota et al., 2001; Wilkins and Marlowe, 2006; Heyer et al., 2012). Around the time when farming spread and social structures changed, many regions of the world show a sharp reduction in Y-chromosome (paternal) diversity and the expansion of particular paternal lineages (Karmin et al., 2015; Poznik et al., 2016). These patterns have been attributed to competition or demographic structure among patrilineal kin groups, or to increased variance in reproductive success among paternal lineages (Zeng et al., 2018; Guyon et al., 2024). Ancient-genome studies comparing sex-chromosomal information with autosomal ancestry have likewise revealed both sex-unbiased and male-biased events during the Neolithic and Bronze Age of Europe (Goldberg et al., 2017; Saag et al., 2017; Olalde et al., 2019). Sex bias can appear not only in the incoming population but also in the local one: during the Middle and Late Neolithic in parts of central and western Europe, the resurgence of hunter–gatherer ancestry among farming populations was male-biased (Mathieson et al., 2018). In East Asia, a male-driven genetic contribution has been inferred during the farming-related subsistence transition in the West Liao River region of northern China (Nakagome and Cooke, 2024). Sex-biased admixture is therefore a general question in reconstructing how changes in subsistence and the movement of people shaped human populations.

In the Japanese archipelago, hints that the admixture between the Jomon hunter– gatherers and the continental farmers was sex-specific have come from two directions: skeletal morphology and uniparental genetic markers (Matsumura, 1990; Okazaki, 2005; Osada and Kawai, 2021). Several regional morphological studies have been interpreted as suggesting sex differences in continental-related affinities, although the evidence derives from different periods, regions, traits, and burial assemblages. In the Kofun-period populations of eastern Japan, the female dentition is closer to that of the Jomon than the male dentition is, suggesting a stronger morphological affinity to the continental migrants among males than among females (Matsumura, 1990). In the Kinki region, the pronounced sexual dimorphism in tooth-crown size has similarly been interpreted as consistent with a male-biased influx of large-toothed migrants (Okazaki, 2005). At the Kuma-Nishioda Yayoi site in northern Kyushu, the subadult skeletons likewise show a male-skewed trend (10 males versus 3 females; Okazaki, 2005). The sex ratio of a burial assemblage is, however, affected by burial selection and mortality structure, and need not directly reflect the sex ratio of the living population or of the incoming group; these remain inferences from particular regions, periods, and burial assemblages.

Genetic lines of evidence point more specifically to a sex difference. The contribution of Jomon ancestry is not consistent among the autosomes, the Y chromosome, and mtDNA: the Jomon-associated Y-chromosome haplogroup D1a2a (D-M55) reaches about 35% in present-day mainland Japanese males and about 81% in the Ainu (Tajima et al., 2004; Hammer et al., 2006; Shi et al., 2008; Sato et al., 2014; Watanabe et al., 2019), whereas both mitochondrial and autosomal Jomon ancestry are found at much lower frequencies. On the maternal side, the putatively Jomon-derived mitochondrial haplogroups N9b and M7a occur in present-day mainland Japanese at frequencies well below that of the Y-chromosomal D1a2a (Tanaka et al., 2004; Adachi et al., 2009); assuming that these haplogroups reached about 70% in the Jomon, the maternal Jomon contribution has been estimated at only around 15% (Osada and Kawai, 2021). Jomon autosomal ancestry is likewise low, with genome-wide estimates of about 10–12% (Kanzawa-Kiriyama et al., 2017, 2019; Gakuhari et al., 2020; Cooke et al., 2021; Watanabe and Ohashi, 2023). Osada and Kawai (2021) named this imbalance the “admixture paradox.” The most intuitive explanation is that Jomon males contributed relatively more than Jomon females at admixture (asymmetric mating between Jomon males and continental migrant females). This “Jomon male-biased” hypothesis, however, has never been tested quantitatively, because the frequency of a single uniparental lineage is shaped not only by the sex ratio at admixture but also by later drift, founder effects, and selection, and therefore cannot by itself reveal that ratio. A high D1a2a frequency could reflect either a male-biased Jomon contribution or these sex-neutral processes, and the uniparental data alone cannot distinguish between them.

Resolving the admixture paradox requires evidence from the whole genome rather than from uniparental markers alone. Because females carry two X chromosomes whereas males carry one, the X chromosome reflects the female side of population history more strongly than the autosomes. Under an equal sex ratio, the ratio of X-chromosomal to autosomal effective population size is expected to be 3/4, and departures from this value can reveal sex-specific demographic processes (Hammer et al., 2008; Keinan et al., 2009). For example, Goldberg et al. (2017) applied this framework to prehistoric Europe and reported that the Neolithic expansion of Anatolian farmers was largely sex-balanced, whereas the later Bronze Age migration from the Pontic–Caspian Steppe was strongly male-biased. Here, we extend this framework by estimating ancestry-specific allele frequencies from local-ancestry markers, allowing the two ancestral components of the modern Japanese genome to be analysed separately. We directly test the long-standing hypothesis that the admixture paradox reflects a male-biased Jomon contribution. Contrary to this expectation, we find that it is better explained by a female-biased contribution of continental East Asian-related ancestry—the first genome-wide test of sex bias in the formation of the modern Japanese population.

## Materials and Methods

### Modern and ancient genomes

For the modern Japanese, we used phased genome data of about 9,290 individuals from the National Center Biobank Network (NCBN; Kawai et al., 2023). For the X chromosome we used female individuals only (4,722 individuals). For the ancient Jomon, we used 42 previously reported genomes for the autosomal analysis (Kanzawa-Kiriyama et al., 2019; Gakuhari et al., 2020; McColl et al., 2018; Cooke et al., 2021; Wang et al., 2021; Watanabe et al., in press; Supplementary Table 1). The X-chromosomal analysis requires the sex of each individual to set the haploid (male) or diploid (female) ploidy, so we restricted it to the 35 individuals with a reliably assigned sex: for the genomes reported by Watanabe et al. (in press) we required the genetic sex to agree with the morphological (anthropological) sex, and for the other previously reported genomes we used the genetic sex (21 females, 14 males). Genotypes were imputed separately for the autosomes and the X chromosome with GLIMPSE2 (Rubinacci et al., 2023), using an NCBN + 1000 Genomes reference panel of 11,772 individuals (both sexes) for the autosomes and a female-only subset of 5,979 individuals for the X chromosome. As proxies of the continental East Asian migrant source, we used present-day Han Chinese (CHB, 103 individuals) and Koreans (1000 Genomes Project Consortium, 2015; Kim et al., 2020). Koreans are used here because Yayoi-period genomes show the highest genetic similarity to present-day Koreans among modern populations and can be modelled as a mixture of Jomon and Korean-related ancestry (Kim et al., 2025, 2026).

### Ancestry-specific allele frequencies in modern Japanese genomes

To detect sex bias within each ancestry component, we needed the allele frequencies of the Jomon and continental ancestries as they are carried by present-day Japanese, not those of the ancient Jomon or present-day continental source populations. Because the sex ratio at admixture is imprinted on the ancestry that was passed on to, and now segregates within, the admixed population, the informative quantities are the allele frequencies of the Jomon-derived and continental-derived portions of modern Japanese genomes. We estimated these directly, as follows.

Jomon-derived variants were identified with the Ancestry Marker Index (AMI), a method we developed previously (Watanabe and Ohashi, 2023). Because the Jomon lineage diverged deeply from other East Asian populations, the modern Japanese genome retains Jomon-specific variants that are not seen in other present-day East Asians; AMI extracts modern-Japanese-specific variants that are in strong linkage disequilibrium with one another and thereby identifies variants of Jomon origin. AMI was applied to 1,972,072 autosomal and 101,318 X-chromosomal SNPs. The custom script that computes AMI and identifies Jomon-derived variants will be released on Zenodo, with the DOI given under Data and Code availability.

We then classified each phased haplotype by ancestry. For each focal SNP, we searched a surrounding window for Jomon-derived variants. The criterion was not simply whether a Jomon-derived variant lay nearby, but whether the same haplotype carried a Jomon-derived allele: a haplotype carrying a Jomon-derived allele within the window was labelled “Jomon-derived,” and a haplotype that did not was labelled “continental East Asian-derived.” Thus, when only one of an individual’s two haplotypes carried a Jomon-derived allele, the two haplotypes were treated as Jomon-derived and continental East Asian-derived, respectively, even around the same focal SNP.

At each focal SNP we computed three allele frequencies: the frequency within Jomon-derived haplotypes (the estimated allele frequency of the Jomon ancestry component), the frequency within continental East Asian-derived haplotypes (that of the continental East Asian ancestry component), and the frequency across all modern Japanese haplotypes. These were computed on both the autosomes (6,992,136 SNPs) and the X chromosome (221,774 SNPs). The estimation was implemented in Rust (jomon_freq); the version used for the analyses and a faster version that gives identical output will be released on Zenodo, with the DOI given under Data and Code availability.

The detection of Jomon-derived variants depends on an AMI threshold, which affects the accuracy of the estimated allele frequencies. We compared two thresholds (T1, AMI > 1; T20, AMI > 20) by comparing the estimated ancestry-specific frequencies with observed frequencies—the estimated frequency in the Jomon-related ancestry against the ancient Jomon frequency, and the estimated frequency in the continental East Asian-related ancestry against the Korean frequency. Across SNPs, the median squared difference was essentially the same for the two thresholds (Jomon-related ancestry component, 0.0064 for T1 vs 0.0069 for T20; continental East Asian-related ancestry component, 0.00044 for both), so for a typical SNP the threshold made little difference. The mean squared difference, in contrast, differed between thresholds (Jomon-related ancestry component, 0.030 for T1 vs 0.047 for T20; continental East Asian-related ancestry component, 0.0058 for T1 vs 0.0021 for T20), indicating that the difference was driven by a minority of SNPs with large discrepancies—those at which the ancestry-specific frequency is estimated least reliably. For each component we adopted the threshold that yielded fewer such large discrepancies, namely T1 for the Jomon-related ancestry component and T20 for the continental East Asian-related ancestry component. Because Q is a ratio between the X chromosome and the autosomes, any threshold dependence common to both chromosomes is partly cancelled.

### Two tests for sex-biased admixture

We tested for sex-biased admixture with two complementary methods.

#### Method 1: chromosome-wise f4-ratio

Using the f4-ratio (admixtools2; Patterson et al., 2012), we estimated the continental East Asian-related ancestry proportion α = f4(YRI, CDX; Japanese, Jomon) / f4(YRI, CDX; CHB, Jomon), separately for the autosomes and for each chromosome including the X, and report the Jomon-related proportion as 1 − α. Here Yoruba (YRI) is the outgroup, Dai (CDX) and Han Chinese (CHB) are present-day continental East Asian references, and the ancient Jomon serve as the reference in the form of their directly observed genotypes—not the Jomon-ancestry allele frequencies that we reconstruct within modern Japanese for the Q statistic (Method 2). If admixture was not sex-biased, the Jomon-related proportion on the X should agree with that on the autosomes; if it was sex-biased, the two should differ, because the X reflects the female side more strongly.

#### Method 2: the Q statistic

Under an equal sex ratio the X-chromosomal effective population size is 3/4 of the autosomal one. Following Keinan et al. (2009), we used

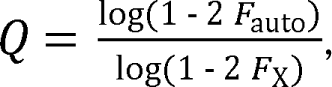

where F_auto_ and F_X_ are the F_ST_ values on the autosomes and the X chromosome, and the neutral expectation under an equal sex ratio is 0.75. The differentiation statistic we use is the ascertainment-conditioned, ratio-of-sums measure of Keinan et al. (2009), for which the expected value is 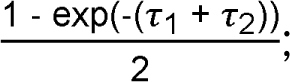 consequently -log(1 – 2*F*) estimates the total genetic drift τ□ + τ□accumulated along the two lineages, and Q is the autosome-to-X ratio of this drift. Each Q compares an ancestral source population (population 1) with the same-ancestry component reconstructed within present-day Japanese (population 2): for the Jomon component, from F_ST_ between the ancient Jomon genomes (population 1) and the Jomon-related ancestry component frequencies estimated within modern Japanese (population 2); for the continental East Asian-related ancestry component, from F_ST_ between continental East Asians (Han Chinese or Koreans; population 1) and the continental East Asian-related ancestry component frequencies estimated within modern Japanese (population 2).

The neutral expectation of 0.75 holds under the four assumptions of Keinan et al. (2009): (1) the focal SNPs are ascertained in population 1 (the source, or reference, population)—that is, for each component we use only sites that are polymorphic in population 1: the ancient Jomon for the Jomon-related ancestry component, and present-day Koreans (or Han Chinese) for the continental East Asian-related ancestry component; (2) population 1 has been approximately panmictic and of constant size since divergence; (3) population 2 has been panmictic since divergence, though not necessarily constant in size. In our application, population 2 is reconstructed from only the subset of source chromosomes transmitted through the admixture, so it inevitably carries a founder effect in addition to subsequent drift. Assumption (3) accommodates this because it requires population 2 to be panmictic but not constant in size; and (4) no substantial gene flow has occurred between population 1 and population 2 since divergence. We assume that later migration from the source populations into the Japanese population was limited, and we separately use the local-ancestry classification to minimize cross-ancestry contamination of the reconstructed components. In the specific demographic scenarios simulated by Keinan et al. (2009), 10% post-split gene flow changed Q only modestly, from 3/4 to 0.747. Two consequences of assumption (3) should be kept in mind. First, population 2 is allowed to experience both the founder effect introduced at admixture and subsequent genetic drift, which increase the absolute amount of drift accumulated on both the autosomes and the X chromosome. Because Q is defined as the ratio of these two drift estimates, the overall magnitude of any sex-neutral drift cancels, making Q primarily sensitive to differences between the autosomes and the X chromosome arising from sex-specific demography— most notably unequal numbers of male and female founder chromosomes at admixture— rather than to the total amount of drift. Q therefore provides a robust test for departure from the neutral expectation of 0.75, and this detection is the primary result reported here. Second, interpreting the magnitude of such a departure as the founder sex ratio is a separate inference. It does not follow from assumptions (1)–(4) alone, but requires the additional assumptions described below (Translating Q into a sex ratio).

Statistical uncertainty was assessed by a block jackknife. For the autosomal blocks we removed each block in turn, recomputed F_auto_ (holding F_X_ fixed), and recomputed Q; for the X-chromosomal blocks we did the same with F_X_. We computed the jackknife variance separately for the autosomes and the X, and summed the two to obtain SE(Q). Departure from the neutral value was assessed by a two-sided z test (asymptotic normal approximation), z = (Q − 0.75) / SE(Q). We also directly tested the difference between the two components, Δ = Q_farming_ − Q_HG_, by removing the same block from both components in each iteration and recomputing both values, so that the covariance between the components is captured. We computed Q over a grid of minor-allele-frequency (MAF) thresholds (0, 0.05, 0.1, 0.2, and 0.3) and physical block sizes (autosomes: 10 Mb, 1 Mb, or 100 kb; X chromosome: 10 Mb, 1 Mb, 100 kb, or 10 kb, with the autosomal block always at least as large as the X block). Figure 2A and the values quoted in the text correspond to a single representative condition (autosome block 1 Mb, X block 100 kb, MAF 0.2), and the full grid is summarized in Figure 2B (Supplementary Table 2). We also checked robustness against the window size and the choice of continental East Asian reference population (Han Chinese or Koreans).

### Translating Q into a sex ratio

The main result of this study is the detection of sex bias by Q; the conversion to a sex ratio below is a secondary step, used mainly to provide inputs for the D1a2a simulation. To translate Q into a founder-model-equivalent sex ratio, we adopt a single-pulse founder model. This requires three additional assumptions: (i) admixture is approximated by a single founder event without substantial later source-specific migration; (ii) subsequent drift and selection did not materially alter the X-to-autosome ratio; and (iii) the reference-population allele frequencies approximate those of the source populations at admixture. Under the founder model, we map the F_ST_-based drift ratio directly to the ratio of founder chromosome copies. With N_f_ females and N_m_ males, the number of X copies is 2N_f_ + N_m_ and the number of autosomal copies is 2(N_f_ + N_m_), so that, with 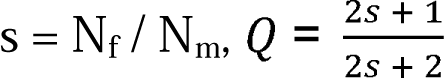 (giving Q = 0.75 at s = 1). Solving for s gives 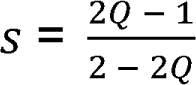 We denote the two component sex ratios s_HG_ and s_farming_. Because the relationship between Q and the founder-model-equivalent sex ratio is highly nonlinear near Q = 1, we interpret Q primarily as a statistic for detecting and comparing sex bias rather than for estimating a precise female-to-male ratio.

### Monte Carlo simulation of D1a2a frequency

We asked whether the inferred sex ratios alone can account for the observed high frequency (∼35%) of the Y-chromosome haplogroup D1a2a in present-day mainland Japanese males. From Method 2, the male fraction of each component’s contributors is 2(1 − Q). With the autosomal Jomon-related ancestry proportion a, the Jomon-related male contribution is p_J_ = a · 2(1 − Q_J_) and the continental East Asian-related male contribution is p_C_ = (1 − a) · 2(1 − Q_C_); the paternal Jomon fraction, taken as the expected D1a2a frequency, is D = p_J_ / (p_J_ + p_C_), assuming that all Jomon Y chromosomes are D1a2a. In each of 10□ iterations we drew Q_J_ = 0.777 ± 0.028, Q_C_ = 0.878 ± 0.028 (reference-population ascertainment; autosome block 1 Mb, X block 100 kb, MAF 0.2), and a = 0.12 ± 0.013 (Kanzawa-Kiriyama et al., 2019; Gakuhari et al., 2020; Cooke et al., 2021) from normal distributions, drawing Q directly so that the non-linearity of the Q-to-s transformation is carried correctly, and compared the resulting distribution of D with the observed value.

## Results

### The chromosome-wise f4-ratio gives a directional signal

The f4-ratio gave a Jomon-related ancestry proportion (1 − α) of about 0.10–0.20 on the autosomes (mean ∼0.14; Figure 1). On the X chromosome the point estimate was lower, near zero, but with a large standard error that could not reject zero; this does not mean that the Jomon-related ancestry proportion is literally zero, but indicates, in direction, a smaller Jomon-related contribution on the X. Chromosome 19 was a similar outlier with a large standard error. This chromosome-wise analysis therefore provides only a directional indication; the statistically robust evidence for sex bias comes from the Q statistic (Method 2).

**Figure 1.**
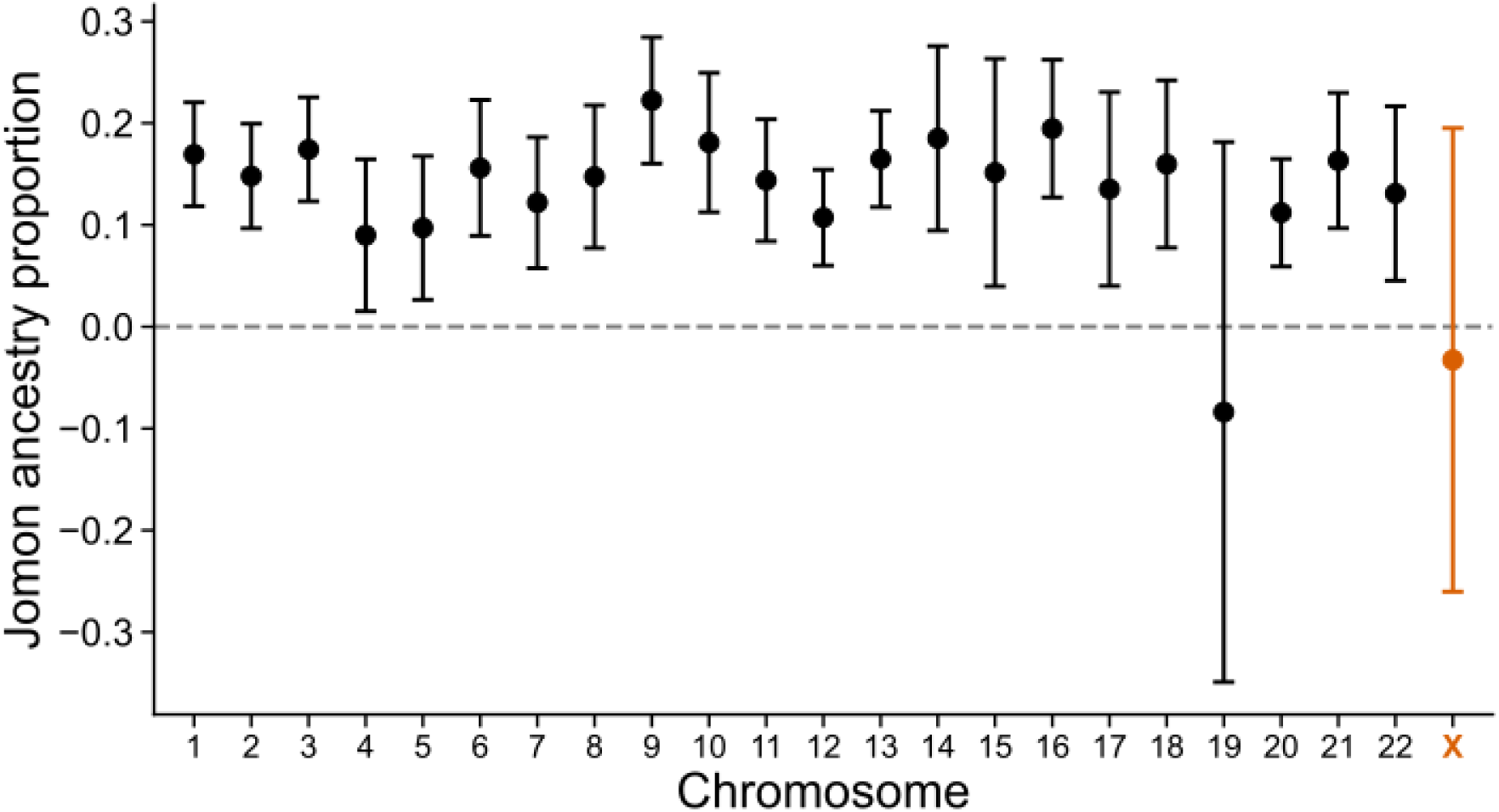
Jomon-related ancestry proportion by chromosome in the modern Japanese genome. Using the f4-ratio with YRI, CDX, present-day Han Chinese (CHB), and ancient Jomon as reference populations, the continental East Asian-related ancestry proportion α was estimated for each chromosome (autosomes 1–22 and the X), and the Jomon-related ancestry proportion is shown as 1 − α. Points are estimates, error bars are 95% confidence intervals, and the dashed line marks zero. The autosomes (black) are distributed around a mean of about 0.14, whereas the X chromosome (orange) has a low point estimate but a large standard error that cannot reject zero. Chromosome 19 is a similar outlier with a large standard error.

### The continental East Asian-related contribution is female-biased, the Jomon-related contribution is not

Our primary interest was the relative difference in sex bias between the two components. We therefore first tested the difference between the two components: Δ = Q_farming_ − Q_HG_ = 0.10 (Figure 2A, SE = 0.039, z = 2.6, P = 9.1 × 10 ³; joint block jackknife; autosome block 1 Mb, X block 100 kb, MAF 0.2), showing that the continental East Asian-related contribution was significantly more female-biased than the Jomon-related contribution. We also compared each component with the neutral value of 0.75. The Jomon-related ancestry component gave Q = 0.78 (P = 0.34), not significantly different from neutrality; thus providing no support for the Jomon male-biased hypothesis. The continental East Asian-related ancestry component gave Q = 0.88 (P = 4.2 × 10□ □), significantly above the neutral value, indicating a relatively larger contribution on the X and thus a female-biased contribution. The signal was robust to the MAF threshold, the block size, and the choice of continental East Asian reference population (Han Chinese or Koreans): across all these conditions the Jomon-related Q stayed below the continental East Asian-related Q, so that the continental East Asian-related female bias was consistent (Figure 2B and Supplementary Table 2). Methods 1 and 2 therefore provide a consistent picture: the continental East Asian-related contribution was more female-biased than the Jomon-related contribution, with the Q of the former significantly above the sex-balanced expectation of 0.75, whereas the Q of the latter showed no detectable departure from that expectation.

**Figure 2.**
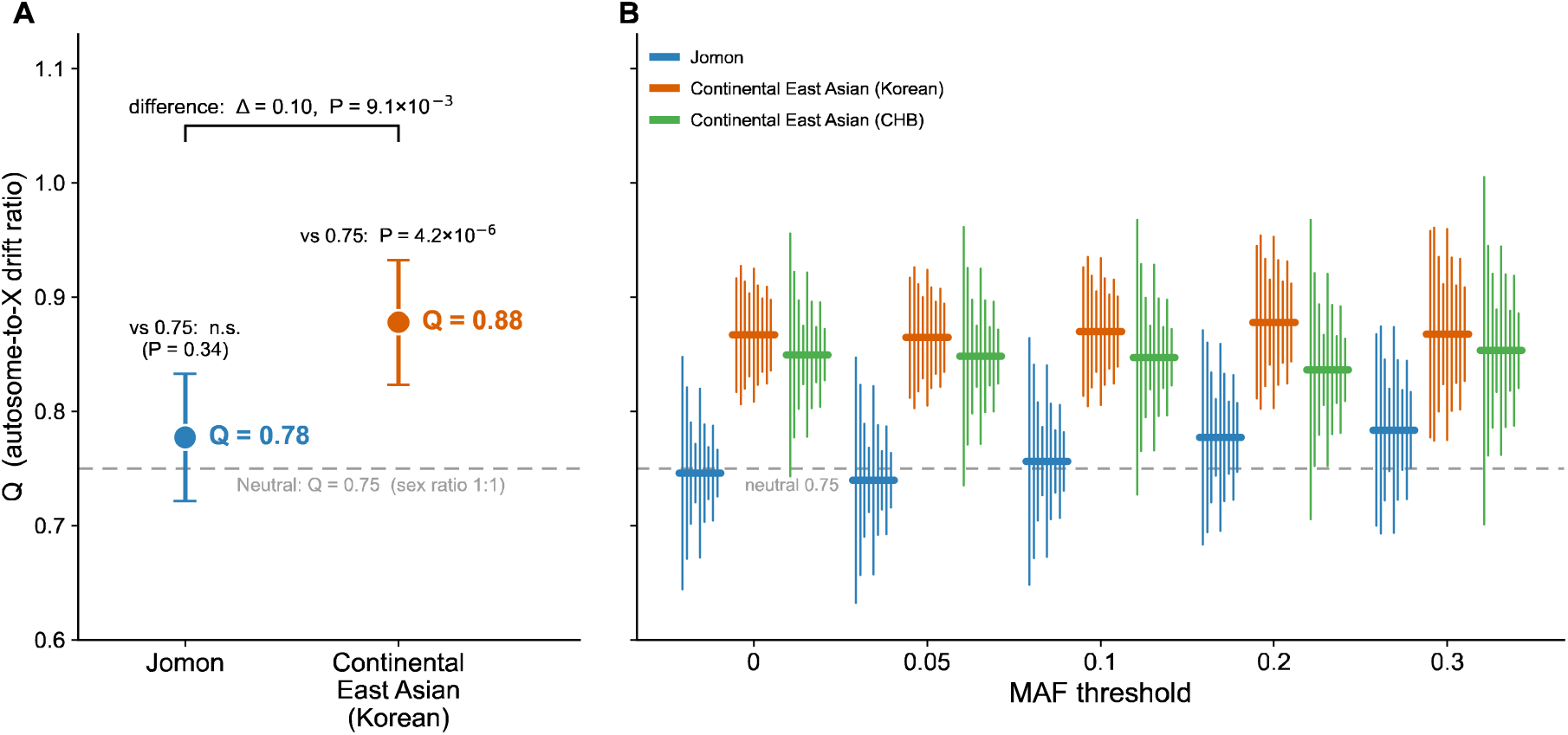
(A) Autosome-to-X drift ratio Q for the Jomon-related and continental East Asian-related ancestry components in the modern Japanese genome. For each component, Q = log(1 − 2 F_auto_) / log(1 − 2 F_X_), where F_auto_ and F_X_ are the F_ST_ values on the autosomes and the X chromosome. Points are estimates and error bars are 95% confidence intervals (estimated by a block jackknife that recomputes Q with each block removed, computing the variance separately for the autosomes and the X and summing them). The dashed line is the neutral expectation of 0.75 under an equal sex ratio, and the P values are two-sided tests against 0.75. The bracket at the top shows the direct test of the difference between the two components (joint block jackknife), the primary test here: Δ = Q_farming_ − Q_HG_ = 0.10, P = 9.1 × 10 ³. Relative to the neutral value, the Jomon-related component does not differ significantly from 0.75 (Q = 0.78, P = 0.34), whereas the continental East Asian-related component is significantly higher (Q = 0.88, P = 4.2 × 10□ □), indicating a female-biased genetic contribution of the continental East Asian-related ancestry. The continental East Asian reference population is Koreans. (B) Robustness of the estimated Q to the MAF threshold, the block size, and the continental East Asian reference population (Koreans and present-day Han Chinese, CHB). Thick horizontal bars are the point estimates under each condition, and thin vertical lines are the 95% confidence intervals for each block size. Across all MAF thresholds, block sizes, and both reference populations, the Jomon-related Q lies below the continental East Asian-related Q, so that the continental East Asian-related female bias (Q_farming_ > Q_HG_) is consistent.

### The inferred sex ratios can account for the high D1a2a frequency

Converting the component Q values into female-to-male sex ratios s gave point estimates of s_HG_ ≈ 1.2 (about 1 male to 1.2 females) for the Jomon-related ancestry component and s_farming_ ≈ 3.1 (about 1 male to 3.1 females) for the continental East Asian-related ancestry component (reported as point estimates only). Given these inferred sex ratios, the expected frequency of D1a2a had a median of 20% (95% CI 13–33%), and the observed value of about 35% lay just above this distribution (Figure 3). The inferred sex ratios therefore account for much of the high frequency of D1a2a but somewhat underpredict it.

**Figure 3.**
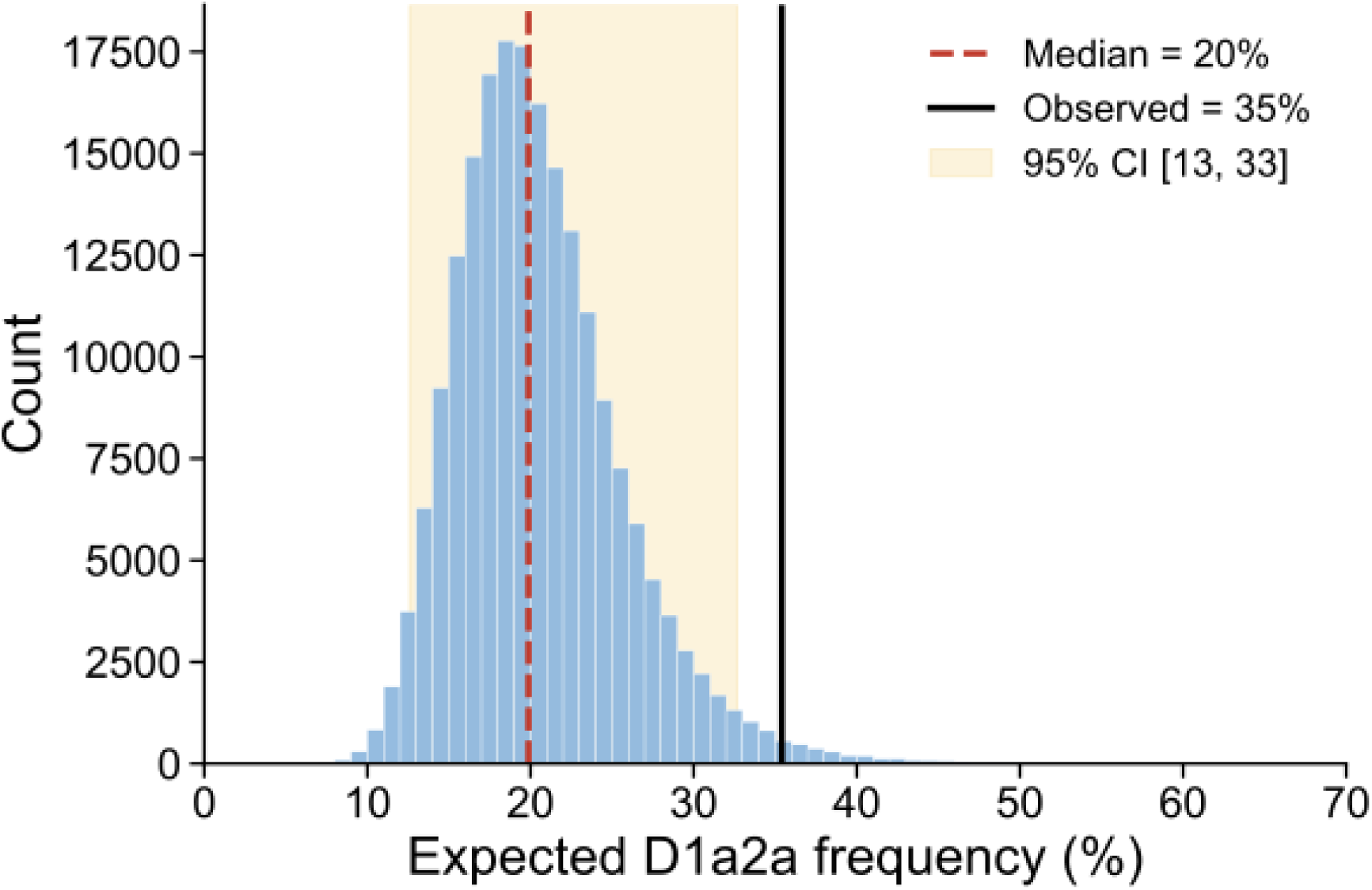
Monte Carlo distribution of the expected frequency of the Y-chromosome haplogroup D1a2a under the inferred admixture sex ratios (10□ iterations). The dashed line is the median of the distribution (20%), the shaded band is the 95% confidence interval (13–33%), and the solid line is the observed frequency in present-day mainland Japanese males (∼35%). The observed value lies just above the distribution, showing that the inferred admixture sex ratios account for much of the high frequency of D1a2a while somewhat underpredicting it.

## Discussion

### Reinterpreting the admixture paradox

The principal finding of this study is that the continental East Asian-related ancestry component shows a significantly higher Q than the Jomon-related ancestry component (Δ = Q_farming_ − Q_HG_ = 0.10, P = 9.1 × 10 ³), demonstrating that sex bias differed between the two ancestral components of the modern Japanese genome. Relative to the neutral value of 0.75, the Jomon-related ancestry component is indistinguishable from neutrality (Q = 0.78), whereas the continental East Asian-related ancestry component is significantly female-biased (Q = 0.88). This does not support the conventional, implicit interpretation of the admixture paradox as a male-biased Jomon contribution; instead, the ancestry structure of the modern Japanese genome was shaped mainly by a female-biased contribution of continental East Asian-related ancestry—that is, the incoming farmers.

A female-biased continental East Asian-related ancestry contribution is also consistent with the high frequency of D1a2a on the Y chromosome. If continental East Asian females contributed disproportionately, then, relative to them, Jomon males are more likely to have left paternal lineages, which is compatible with the high frequency of Jomon-associated D1a2a on the Y chromosome. In our simulation, the observed D1a2a frequency (∼35%) lay just above the distribution expected from the inferred sex ratios (median 20%, 95% CI 13–33%), so the paradox is largely explained without invoking a male-biased Jomon contribution. This comparison is necessarily approximate because it assumes a single admixture pulse, that all Jomon Y chromosomes were D1a2a, and no subsequent Y-chromosomal drift or selection. Because the model somewhat underpredicts the observed value, the residual points to a continental East Asian female bias even stronger than our point estimate, or to an additional Jomon paternal contribution or post-admixture processes on the Y.

### What might have determined the direction of the sex bias

The direction and degree of sex bias that accompany a shift from foraging to farming, or the movement and mixture of populations, differ among regions and events. In Europe, the Neolithic spread of farming from Anatolia shows no consistent sex bias (Goldberg et al., 2017), whereas the Bronze Age migrations from the Pontic–Caspian steppe show a strong male bias (Goldberg et al., 2017; Saag et al., 2017; Olalde et al., 2019). Sex bias can also appear on the local side of an admixture: during the Middle and Late Neolithic in parts of Europe, the resurgence of hunter–gatherer ancestry among farming populations was male-biased (Mathieson et al., 2018). In East Asia, the farming-related subsistence transition in the West Liao River region of northern China has been interpreted as a male-driven contribution of Yellow-River-related farmers (from a proportion-based X-to-autosome comparison; Nakagome and Cooke, 2024). By contrast, some island admixture histories show a comparable contrast: in Vanuatu the East Asian/Austronesian-related ancestry is female-biased while the Papuan-related ancestry is male-biased (Arauna et al., 2022; Posth et al., 2018; Lipson et al., 2020), and Madagascar shows the same pattern for Austronesian-related versus African/Bantu-related ancestry (Pierron et al., 2017). The female-biased continental farmer contribution found here contrasts with the male-biased or approximately sex-balanced patterns reported in several well-studied cases of admixture accompanying farming dispersals. Together, these comparisons indicate that sex bias accompanying population admixture has no universal direction, and that the Japanese case is an uncommon example in which the incoming farming-related ancestry was female-biased.

The following is more speculative. Because the direction of sex bias in admixture seems to depend on social context rather than on subsistence type alone, we ask what might have produced a female-biased farming-migrant contribution. One candidate mechanism is post-marital residence: in patrilocal societies, women generally move between communities more often than men, potentially producing greater female-mediated gene flow (Oota et al., 2001). Sex-asymmetric genetic patterns consistent with such residence-driven, female-biased gene flow have been reported in eastern North America and Remote Oceania (Bolnick et al., 2006; Arauna et al., 2022). Although any such reconstruction remains speculative, the female-biased farmer contribution could, for instance, have arisen if patrilocal hunter-gatherer communities received incoming farmer women, or if the expansion of farming communities was itself accompanied by female movement. In principle one could restate this symmetrically as a matrilocal continental farming migrant society that received local indigenous hunter-gatherer males; but that scenario predicts a male bias in the indigenous hunter-gatherer component (Q_HG_ < 0.75), which is hard to reconcile with the neutral Jomon hunter-gatherer Q observed here (0.78, P = 0.34). All of these scenarios assume particular residence rules; because foragers are less consistently patrilocal than non-foraging populations and their post-marital residence patterns are diverse (Marlowe, 2004), it is not obvious that the Jomon hunter-gatherers were patrilocal. We have no independent evidence to distinguish these possibilities. Reconstructing the marriage and residence practices of the Jomon-to-Yayoi transition—by combining ancient DNA with archaeological evidence on kinship and residence—would allow this mechanism to be tested directly.

### Interpretation and limitations of Q

The component-specific Q values primarily indicate the direction and relative magnitude of sex bias rather than a unique numerical sex ratio. X-to-autosome contrasts depend on the timing and continuity of admixture, and Q is additionally sensitive to sex-specific changes in population size, population structure, migration, and mismatch between the reference and source populations (Goldberg and Rosenberg, 2015; Ramachandran et al., 2008; Emery et al., 2010). The conversion of Q to a founder-model-equivalent sex ratio therefore applies only under the single-pulse model described above. Moreover, the transformation from Q to the founder-model-equivalent sex ratio is strongly nonlinear: increasingly different sex ratios are compressed into Q values near the extremes (Q → 0.5 for an all-male and Q → 1 for an all-female contribution). For example, founder-model-equivalent sex ratio of 5, 10, and 20 females per male correspond to Q ≈ 0.92, 0.95, and 0.98, respectively, so that markedly different strong female biases map to very similar Q values. Q is therefore well suited for detecting the presence and direction of sex bias but less informative for precisely quantifying its magnitude, which particularly limits the resolution of the female-biased continental East Asian ratio inferred here. Indeed, inverting the chromosome-wise f4-ratio estimates (Figure 1) directly into sex-specific contributions by the approach of Goldberg and Rosenberg (2015) even yields a negative—and hence impossible—female contribution for the Jomon ancestry, reflecting not a real deficit of Jomon females but imprecise estimates that lie outside the range a single-pulse sex-biased admixture can produce. We consequently treat the component sex ratios s_HG_ and s_farming_ as illustrative point estimates only; the primary and robust quantity is Q, and our conclusions rest on Q and its between-component difference Δ rather than on s.

Several features support the relative conclusion. The lack of a detectable departure of the Jomon-related Q from 0.75 is consistent with the absence of a strong global bias affecting both ancestry components in the same direction; the continental East Asian ancestry component remains higher across MAF thresholds, block sizes, ancestry-marker settings, and both Korean and CHB references; and the difference between components is significant in a joint block jackknife. These observations do not exclude component-specific effects arising from differences in reference populations or ancestry reconstruction, but they support the conclusion that the continental farmer contribution was more female-biased than the Jomon hunter-gatherer contribution.

## Conclusion

Using ancestry-derived haplotypes in the modern Japanese genome, we present a framework that detects and compares sex bias separately for each ancestry component. Applying this framework, we show that the long-standing admixture paradox is better explained by a female-biased contribution of continental farming migrants than by a male-biased contribution of the indigenous Jomon hunter-gatherers. More broadly, our findings illustrate how genome-wide analyses are beginning to move beyond reconstructing ancestry alone toward reconstructing prehistoric social organization. As genomic data continue to accumulate and are integrated with archaeology and biological anthropology, they will increasingly illuminate patterns of kinship, mobility, marriage, and interaction, providing a richer understanding of how prehistoric societies shaped human population history.

## Code availability

Custom code used in this study is deposited at Zenodo 10.5281/zenodo.21644608 and includes scripts for detecting Jomon-derived ancestry markers (the Ancestry Marker Index; Watanabe and Ohashi, 2023), the Rust program jomon_freq for estimating ancestry-specific allele frequencies.

## Acknowledgments

This work was supported by JSPS KAKENHI Grant Numbers 26K09483, 25H00999, 24K21381, 23K21990, 23H04840, 23H04838, 23H04836, 23H00009, 22H00421, 22H00020, 21H04779, 21H00337, 21K15175, 19H04526, 18H03590, and 18H03593. Computations were partially performed on the NIG supercomputer at ROIS National Institute of Genetics.

