## Supplementary for "Female-biased continental farmer ancestry contribution to the modern Japanese genome"

**Supplementary Text**

**NCBN Controls WGS Consortium members**

Hatsue Ishibashi-Ueda^1^, Tsutomu Tomita^1^, Michio Noguchi^1^, Ayako Takahashi^1^, Yu-ichi Goto^2^, Sumiko Yoshida^3^, Kotaro Hattori^3^, Ryo Matsumura^3^, Aritoshi Iida^4^, Yutaka Maruoka^5^, Hiroyuki Gatanaga^6^, Akihiko Shimomura^5^, Masaya Sugiyama^7^, Satoshi Suzuki^5^, Kengo Miyo^8^, Yoichi Matsubara^9^, Akihiro Umezawa^10^, Kenichiro Hata^11^, Tadashi Kaname^12^, Kouichi Ozaki^13^, Haruhiko Tokuda^13^, Hiroshi Watanabe^13^, Shumpei Niida^13^, Eisei Noiri^14^, Koji Kitajima^14^, Yosuke Omae^14,15^, Reiko Miyahara^14^, Hideyuki Shimanuki^14^, Yosuke Kawai^15^, and Katsushi Tokunaga^14,15^.

^1^ NCVC Biobank, National Cerebral and Cardiovascular Center, Suita, Osaka 564-8565, Japan

^2^ Medical Genome Center, National Center of Neurology and Psychiatry, Kodaira, Tokyo 187-8551, Japan

^3^ Department of Bioresources, Medical Genome Center, National Center of Neurology and Psychiatry, Kodaira, Tokyo 187-8551, Japan

^4^ Department of Clinical Genome Analysis, Medical Genome Center, National Center of Neurology and Psychiatry, Kodaira, Tokyo 187-8551, Japan

^5^ NCGM Biobank, National Center for Global Health and Medicine, Shinjuku-ku, Tokyo 162-8655, Japan

^6^ AIDS Clinical Center, National Center for Global Health and Medicine, Shinjuku-ku, Tokyo 162-8655, Japan

^7^ Department of Viral Pathogenesis and Controls, Research Institute, National Center for Global Health and Medicine, Ichikawa, Chiba 272-8516, Japan

^8^ Center for Medical Informatics and Intelligence, National Center for Global Health and Medicine, Shinjuku-ku, Tokyo 162-8655, Japan

^9^ National Center for Child Health and Development, Setagaya-ku, Tokyo 157-8535, Japan

^10^ Center for Regenerative Medicine, National Center for Child Health and Development, Setagaya-ku, Tokyo 157-8535, Japan

^11^ Department of Maternal-Fetal Biology, National Center for Child Health and Development, Setagaya-ku, Tokyo 157-8535, Japan

^12^ Department of Genome Medicine, National Center for Child Health and Development, Setagaya-ku, Tokyo 157-8535, Japan

^13^ Research Institute, National Center for Geriatrics and Gerontology, Obu, Aichi 474-8511, Japan

^14^ Central Biobank, National Center Biobank Network, Shinjuku-ku, Tokyo 162-8655, Japan

^15^ Genome Medical Science Project (Toyama), Research Institute, National Center for Global Health and Medicine, Shinjuku-ku, Tokyo 162-8655, Japan

**Supplementary Table legends**

**Supplementary Table 1.** Ancient Jomon individuals analysed in this study (n = 42). All 42 individuals were used for the autosomal analysis. The 35 individuals with an entry in the “Sex (X analysis)” column were also used for the X-chromosome analysis (21 females, 14 males); these are individuals with a reliably assigned sex — for the genomes reported by Watanabe et al. (in press) the genetic sex was required to agree with the morphological (anthropological) sex, whereas for the other previously reported genomes the genetic sex was used. A blank entry denotes an individual used for the autosomes only (KT-2 was excluded from the X analysis because its genetic and anthropological sex disagree; the others lack a determined anthropological sex). Age (YBP): for the 25 individuals reported by Watanabe et al. (in press), uncalibrated radiocarbon age (C14 BP) ± 1 SD; for the other 17 individuals, as reported in the original study. Depth: mean genome coverage (hg38). Accession: DDBJ/DRA or ENA run accessions. All 42 individuals were previously reported (Cooke et al., 2021; Gakuhari et al., 2020; Kanzawa-Kiriyama et al., 2019; Wang et al., 2021, Watanabe et al., in press).

**Supplementary Table 2.** Autosome-to-X drift ratio Q underlying Figure 2B. Q is shown for the indigenous hunter-gatherer component and for the continental farming-migrant component (with Korean and Han Chinese, CHB, references), computed under reference-population ascertainment (ascertain = ref) with the uncorrected (raw) Keinan/Hudson ratio-of-sums F_ST_, across minor-allele-frequency (MAF) thresholds and autosome/X block sizes. The 95% confidence interval is Q ± 1.96 × SE, and P is a two-sided test against the neutral value of 0.75. The Q point estimate depends only on the MAF threshold; the block size affects only the standard error (block jackknife).
